# SARS-CoV-2 Spike S1 Receptor Binding Domain undergoes Conformational Change upon Interaction with Low Molecular Weight Heparins

**DOI:** 10.1101/2020.04.29.068486

**Authors:** Courtney J. Mycroft-West, Dunhao Su, Yong Li, Scott E. Guimond, Timothy R. Rudd, Stefano Elli, Gavin Miller, Quentin M. Nunes, Patricia Procter, Antonella Bisio, Nicholas R. Forsyth, Jeremy E. Turnbull, Marco Guerrini, David G. Fernig, Edwin A. Yates, Marcelo A. Lima, Mark A. Skidmore

## Abstract

The dependence of the host on the interaction of hundreds of extracellular proteins with the cell surface glycosaminoglycan heparan sulphate (HS) for the regulation of homeostasis is exploited by many microbial pathogens as a means of adherence and invasion. The closely related polysaccharide heparin, the widely used anticoagulant drug, which is structurally similar to HS and is a common experimental proxy, can be expected to mimic the properties of HS. Heparin prevents infection by a range of viruses when added exogenously, including S-associated coronavirus strain HSR1 and inhibits cellular invasion by SARS-CoV-2. We have previously demonstrated that unfractionated heparin binds to the Spike (S1) protein receptor binding domain, induces a conformational change and have reported the structural features of heparin on which this interaction depends. Furthermore, we have demonstrated that enoxaparin, a low molecular weight clinical anticoagulant, also binds the S1 RBD protein and induces conformational change. Here we expand upon these studies, to a wide range of low molecular weight heparins and demonstrate that they induce a variety of conformational changes in the SARS-CoV-2 RBD. These findings may have further implications for the rapid development of a first-line therapeutic by repurposing low molecular weight heparins, as well as for next-generation, tailor-made, GAG-based antiviral agents, against SARS-CoV-2 and other members of the *Coronaviridae*.

## Introduction

Heparin, the second most widely used drug by weight globally, is formulated as a polydisperse, heterogenous natural product. Unfractionated heparin (UFH), low molecular weight heparins (LMWHs) and heparinoids are clinically approved as anticoagulants / thrombotics with excellent safety, stability, bioavailability and pharmacokinetic profiles. Crucially, heparin and its derivatives, some of which, lacking significant anticoagulant activity ^1^, are an under-exploited antiviral drug class, despite possessing broad-spectrum activity against a multitude of distinct viruses, including *coronaviridae* and SARS-associated coronavirus strains ^2,3^, in addition to flaviviruses ^4,5^, herpes ^6^, influenza ^7^ and HIV ^8,9^.

Traditional drug development processes are slow and ineffective against emerging public health threats such as the current SARS-CoV-2 coronavirus outbreak which makes the repurposing of existing drugs a timely and attractive alternative. Heparin, a well-tolerated anticoagulant pharmaceutical, has been used safely in medicine for over 80 years and alongside its anticoagulant activities, its ability to prevent viral infection, including *coronaviridae*, has been described^1^. Furthermore, the closely related glycosaminoglycan (GAG) member, heparan sulfate (HS), is known to bind CoV surface proteins and to be used by coronavirus for its attachment to target cells^10^. Low molecular weight heparins are derived from unfractionated heparin by different chemical and enzymatic depolymerisation processes. Ultimately, despite being chemically distinct, they do share common chemical features amongst themselves and those from the parent, unfractionated heparin.

Currently, there are no commercially available medicinal products designed to treat and/or prevent infections associated with the new SARS-CoV-2 coronavirus outbreak. Here, we describe preliminary observations suggesting LMWHs have the ability to interact with the SARS-CoV-2 S1 RBD, as previously demonstrated for unfractionated heparin^2^.

## Methods & Materials

### 2.1 Recombinant expression of SARS-CoV-2 S1 RBD

Residues 330-583 of the SARS-CoV-2 Spike Protein (GenBank: MN908947) were cloned upstream of a N-terminal 6XHisTag in the pRSETA expression vector and transformed into SHuffle^®^ T7 Express Competent *E. coli* (NEB, UK). Protein expression was carried out in MagicMedia™ *E. coli* Expression Media (Invitrogen, UK) at 30°C for 24 hrs, 250 rpm. The bacterial pellet was suspended in 5 mL lysis buffer (BugBuster Protein Extraction Reagent, Merck Millipore, UK; containing DNAse) and incubated at room temperature for 30 mins. Protein was purified from inclusion bodies using IMAC chromatography under denaturing conditions. On-column protein refolding was performed by applying a gradient with decreasing concentrations of the denaturing agent (6 - 0 M Urea). After extensive washing, protein was eluted using 20 mM NaH_2_PO_4_, pH 8.0, 300 mM NaCl, 500 mM imidazole. Fractions were pooled and buffer-exchanged to phosphate-buffered saline (PBS; 140 mM NaCl, 5 mM NaH_2_PO_4_, 5 mM Na_2_HPO_4_, pH 7.4; Lonza, UK) using a Sephadex G-25 column (GE Healthcare, UK). Recombinant protein was stored at −20°C until required.

### 2.2 Secondary structure determination of SARS-CoV-2 S1 RBD by circular dichroism spectroscopy

The circular dichroism (CD) spectrum of the SARS-CoV-2 S1 RBD in PBS was recorded using a J-1500 Jasco CD spectrometer (Jasco, UK), Spectral Manager II software (JASCO, UK) and a 0.2 mm pathlength, quartz cuvette (Hellma, USA) scanning at 100 nm.min^−1^ with 1 nm resolution throughout the range λ = 190 - 260 nm. All spectra obtained were the mean of five independent scans, following instrument calibration with camphorsulfonic acid. SARS-CoV-2 S1 RBD was buffer-exchanged (prior to spectral analysis) using a 5 kDa Vivaspin centrifugal filter (Sartorius, Germany) at 12,000 g, thrice and CD spectra were collected using 21 μl of a 0.6 mg.ml^−1^ solution in PBS, pH 7.4. Spectra of UFH (porcine mucosal heparin) and LMWHs were collected in the same buffer at approximately comparable concentrations, since these are disperse materials. Collected data were analysed with Spectral Manager II software prior to processing with GraphPad Prism 7, using second order polynomial smoothing through 21 neighbours. Secondary structural prediction was calculated using the BeStSel analysis server^11^.To ensure that the CD spectral change of SARS-CoV-2 S1 RBD in the presence UFH or LMWH alone did not arise from simply from the addition of the carbohydrate (this class of carbohydrates are known to possess CD spectrum at high concentrations ^12,13^), the difference spectrum was analysed for each carbohydrate in order to verify that the change in the CD spectrum arose from a conformational change following binding to the test carbohydrate.

### 2.3 Surface Plasmon Resonance determination of SARS-CoV-2 S1 RBD binding to unfractionated heparin

Human FGF2 was produced as described by Duchesne *et al*.^16^. Porcine mucosal heparin was biotinylated at the reducing end using hydroxylamine biotin (ThermoFisher, UK) as described by Thakar *et al*. ^17^. Heparin (20 μl of 50 mg.ml^−1^) was reacted with 20 μl hydroxylamine-biotin in 40 μl 300 mM aniline (Sigma-Aldrich, UK) and 40 μl 200 mM acetate pH 6 for 48 h at 37 °C. Free biotin was removed by gel-filtration chromatography on Sephadex G25 (GE LifeSciences, UK).

A P4SPR, multi-channel Surface Plasmon Resonance (SPR) instrument (Affinté Instruments; Montréal, Canada) was employed with a gold sensor chip that was plasma cleaned prior to derivatization. A self-assembled monolayer of mPEG thiol and biotin mPEG was formed by incubating the chip in a 1 mM solution of these reagents at a 99:1 molar ratio in ethanol for 24 hrs^18^. The chip was rinsed with ethanol and placed in the instrument. PBS (1X) was used as the running buffer for the three sensing and a fourth background channel at 500 μl.min^−1^, using an Ismatec pump. Twenty micrograms of streptavidin (Sigma, UK; 1 ml in PBS) were injected over the four sensor channels. Subsequently, biotin-heparin (1 ml) was injected over the three sensing channels.

Binding experiments used PBS with 0.02% Tween 20 (v/v) as the running buffer. The ligand was injected over the three sensing channels, diluted to the concentration indicated (see figures) at 500 μl.min^−1^. Sensor surfaces with bound FGF2 were regenerated by a wash with 2 M NaCl (Fisher Scientific, UK). However, this was found to be ineffectual for SARS-CoV-2 S1 RBD. Partial regeneration of the surface was achieved with 20 mM HCl (VWR, UK) and only 0.25 % (w/v) SDS (VWR, UK) was found to remove the bound protein. After regeneration with 0.25 % (w/v/) SDS, fluidics and surfaces were washed with 20 mM HCl to ensure all traces of the detergent were removed. Background binding to the underlying streptavidin bound to the mPEG/biotin mPEG self-assembled monolayer was determined by injection over the control channel. Responses are reported as the change in plasmon resonance wavelength, in nm and for the three control channels represent their average response.

## Results

### 3.1 Secondary structure determination of SARS-CoV-2 S1 RBD protein by circular dichroism spectroscopy

Circular dichroism (CD) spectroscopy detects changes in protein secondary structure that occur in solution using UV radiation. Upon binding, conformational changes are detected and quantified using spectral deconvolution ^19^. Indeed, SARS-CoV-2 S1 RBD underwent conformational change in the presence of LMWHs (Figures 1 – 6). These observed changes further demonstrate that the SARS-CoV-2 S1 RBD interacts not only with UFH, but also, with its low molecular weight derivatives.

**Figure 1:**
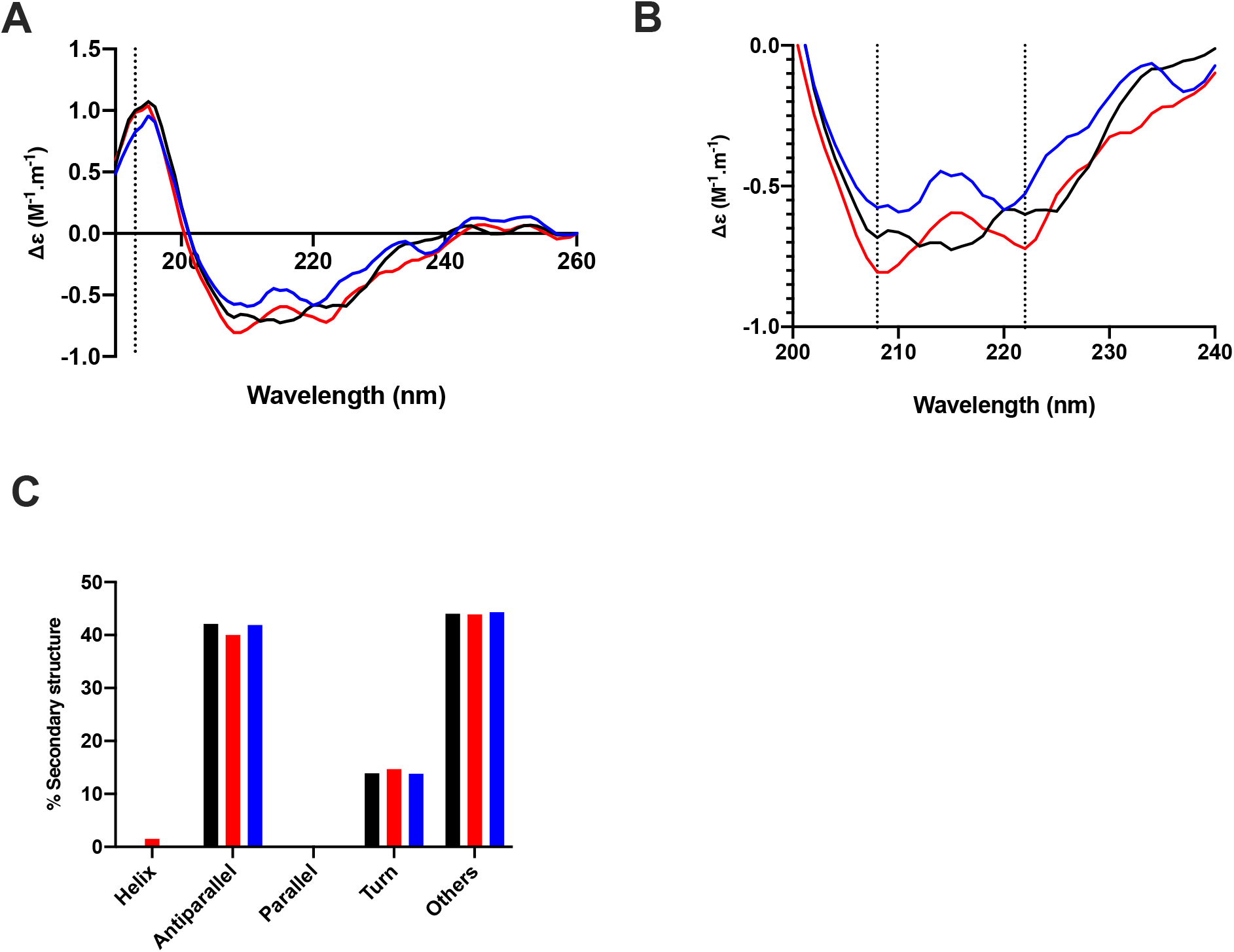
The conformational change of the SARS-CoV-2 S1 RBD observed in the presence of enoxaparin by circular dichroism (CD) spectroscopy. (**A**) Circular dichroism spectra (190 - 260 nm) of nCovS1RBD alone (black solid line), heparin (red solid line) and enoxaparin (blue) in PBS, pH 7.4. The dotted vertical line indicates 193 nm. (**B**) Details of the same spectra expanded between 200 and 240 nm. Vertical dotted lines indicate 222 nm and 208 nm. (**C**) Secondary structure content analysed using BeStSel for nCovS1RBD. α helical secondary structure is characterized by a positive band at ~193 nm and two negative bands at ~208 and ~222 nm (analysis using BeStSel was performed on smoothed data between 190 and 260 nm.

**Figure 2:**
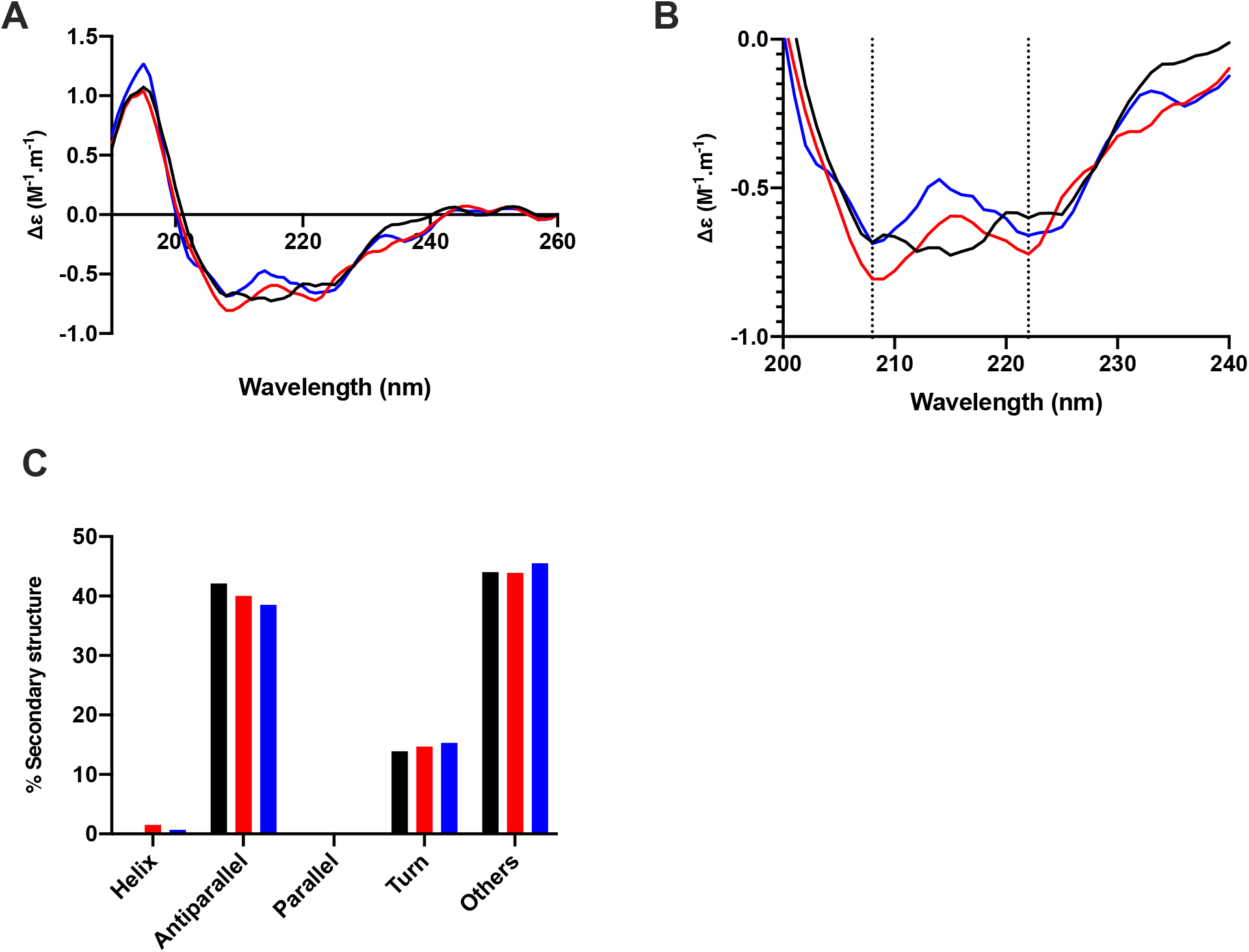
The conformational change of the SARS-CoV-2 S1 RBD observed in the presence of dalteparin by circular dichroism (CD) spectroscopy. (**A**) Circular dichroism spectra (190 - 260 nm) of nCovS1RBD alone (black solid line), heparin (red solid line) and dalteparin (blue) in PBS, pH 7.4. The dotted vertical line indicates 193 nm. (**B**) Details of the same spectra expanded between 200 and 240 nm. Vertical dotted lines indicate 222 nm and 208 nm. (**C**) Secondary structure content analysed using BeStSel for nCovS1RBD. α helical secondary structure is characterized by a positive band at ~193 nm and two negative bands at ~208 and ~222 nm (analysis using BeStSel was performed on smoothed data between 190 and 260 nm.

**Figure 3:**
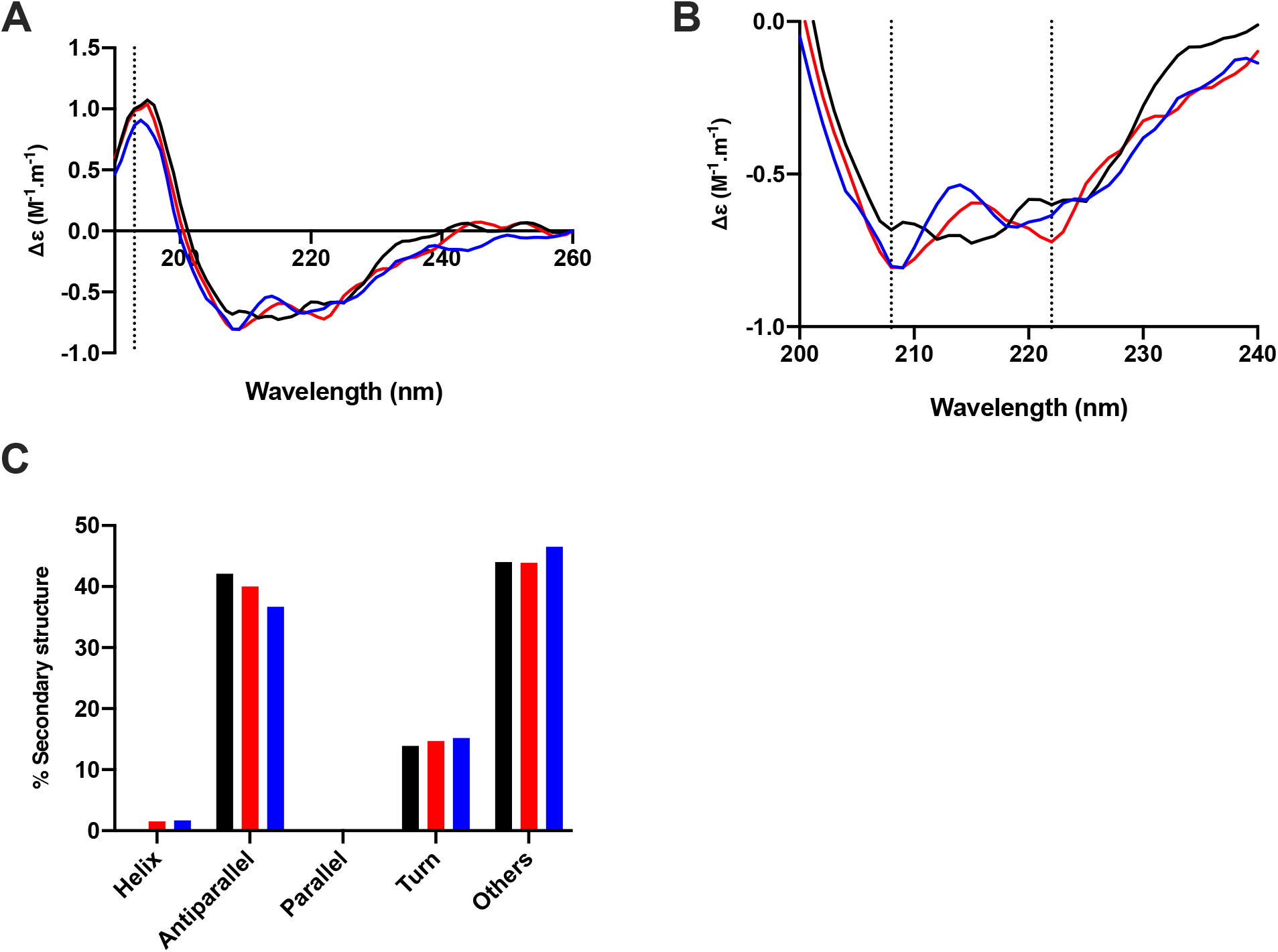
The conformational change of the SARS-CoV-2 S1 RBD observed in the presence of nadroparin by circular dichroism (CD) spectroscopy. (**A**) Circular dichroism spectra (190 - 260 nm) of nCovS1RBD alone (black solid line), heparin (red solid line) and nadroparin (blue) in PBS, pH 7.4. The dotted vertical line indicates 193 nm. (**B**) Details of the same spectra expanded between 200 and 240 nm. Vertical dotted lines indicate 222 nm and 208 nm. (**C**) Secondary structure content analysed using BeStSel for nCovS1RBD. α helical secondary structure is characterized by a positive band at ~193 nm and two negative bands at ~208 and ~222 nm (analysis using BeStSel was performed on smoothed data between 190 and 260 nm.

**Figure 4:**
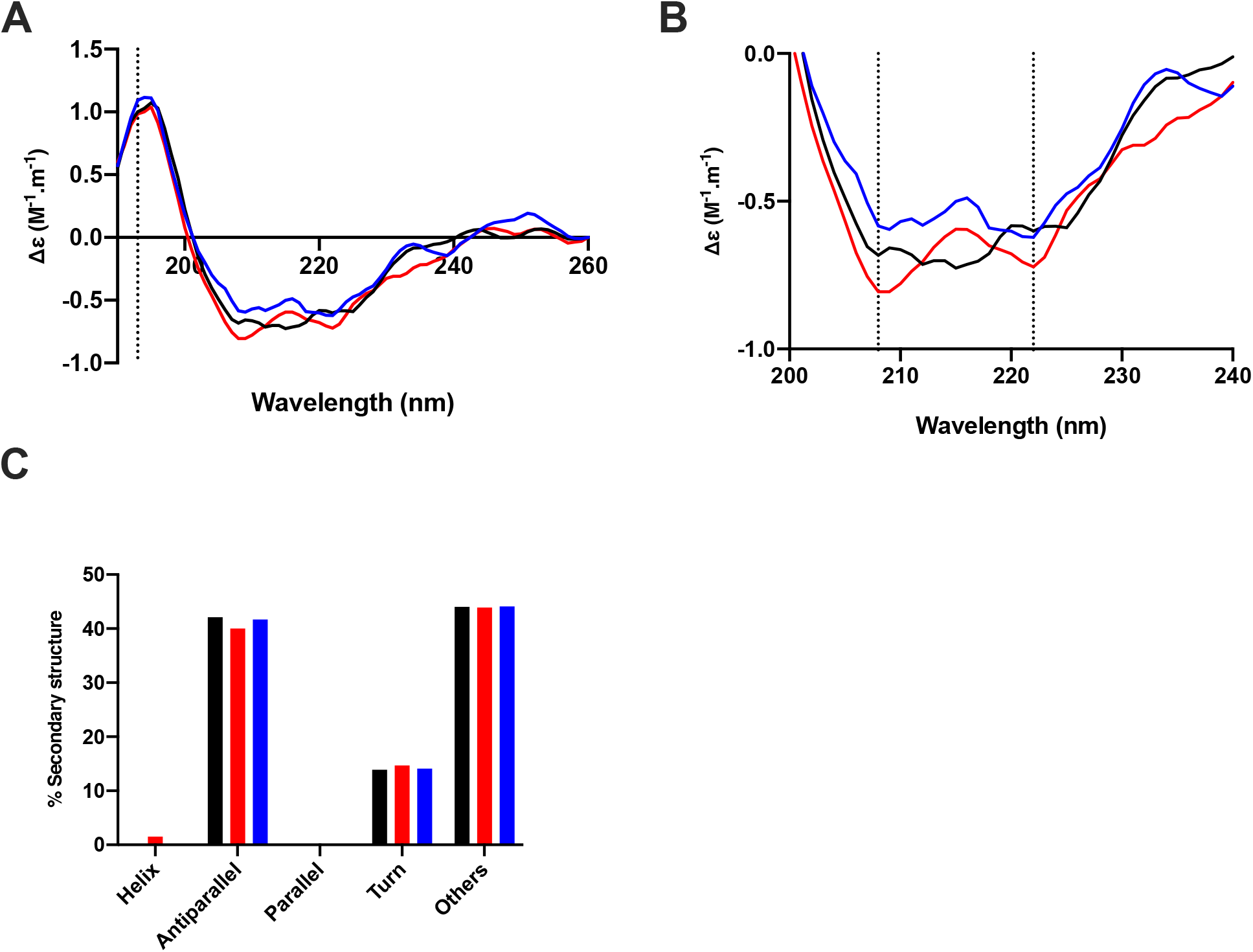
The conformational change of the SARS-CoV-2 S1 RBD observed in the presence of parnaparin by circular dichroism (CD) spectroscopy. (**A**) Circular dichroism spectra (190 - 260 nm) of nCovS1RBD alone (black solid line), heparin (red solid line) and parnaparin (blue) in PBS, pH 7.4. The dotted vertical line indicates 193 nm. (**B**) Details of the same spectra expanded between 200 and 240 nm. Vertical dotted lines indicate 222 nm and 208 nm. (**C**) Secondary structure content analysed using BeStSel for nCovS1RBD. α helical secondary structure is characterized by a positive band at ~193 nm and two negative bands at ~208 and ~222 nm (analysis using BeStSel was performed on smoothed data between 190 and 260 nm.

**Figure 5:**
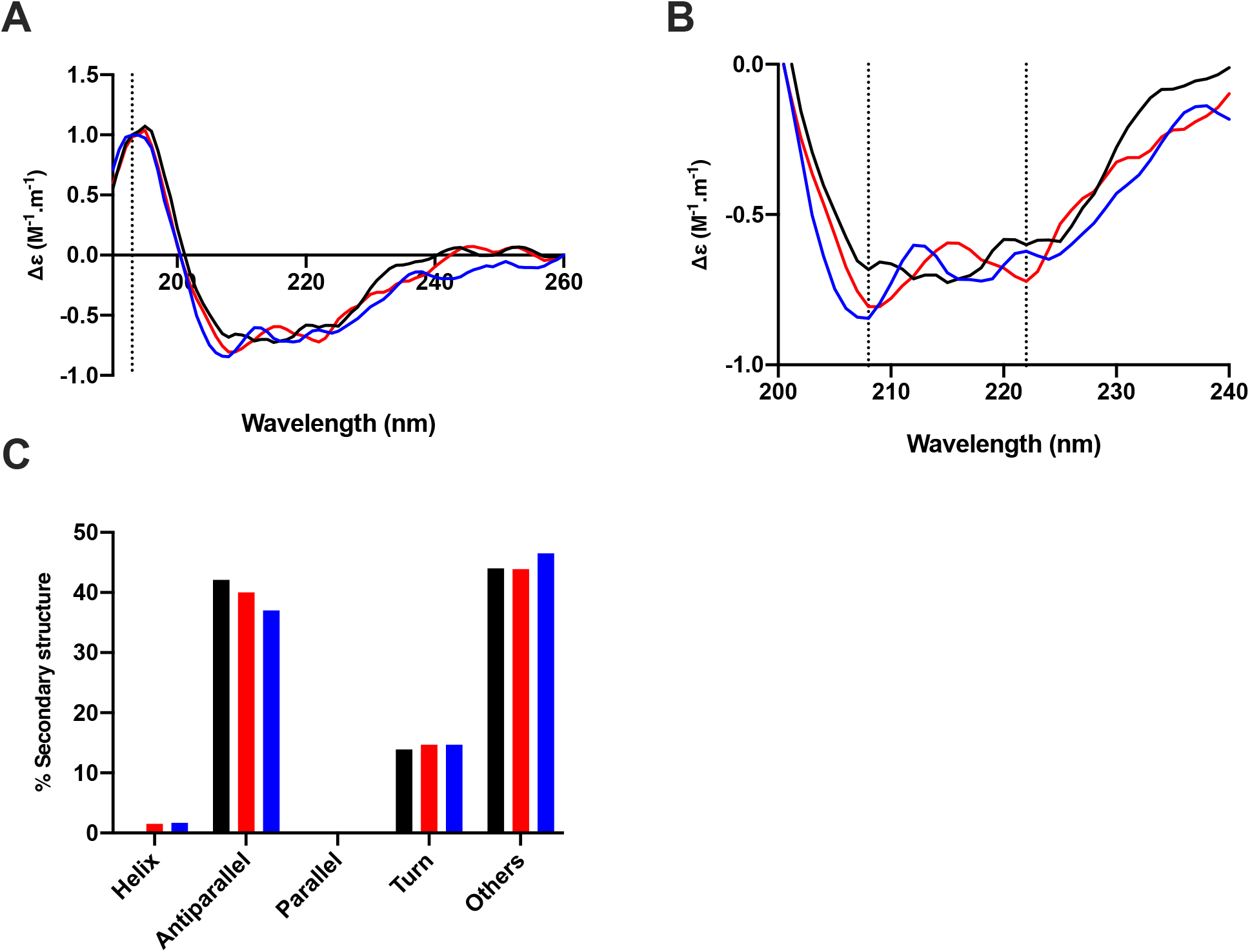
The conformational change of the SARS-CoV-2 S1 RBD observed in the presence of reviparin by circular dichroism (CD) spectroscopy. (**A**) Circular dichroism spectra (190 - 260 nm) of nCovS1RBD alone (black solid line), heparin (red solid line) and reviparin (blue) in PBS, pH 7.4. The dotted vertical line indicates 193 nm. (**B**) Details of the same spectra expanded between 200 and 240 nm. Vertical dotted lines indicate 222 nm and 208 nm. (**C**) Secondary structure content analysed using BeStSel for nCovS1RBD. α helical secondary structure is characterized by a positive band at ~193 nm and two negative bands at ~208 and ~222 nm (analysis using BeStSel was performed on smoothed data between 190 and 260 nm.

**Figure 6:**
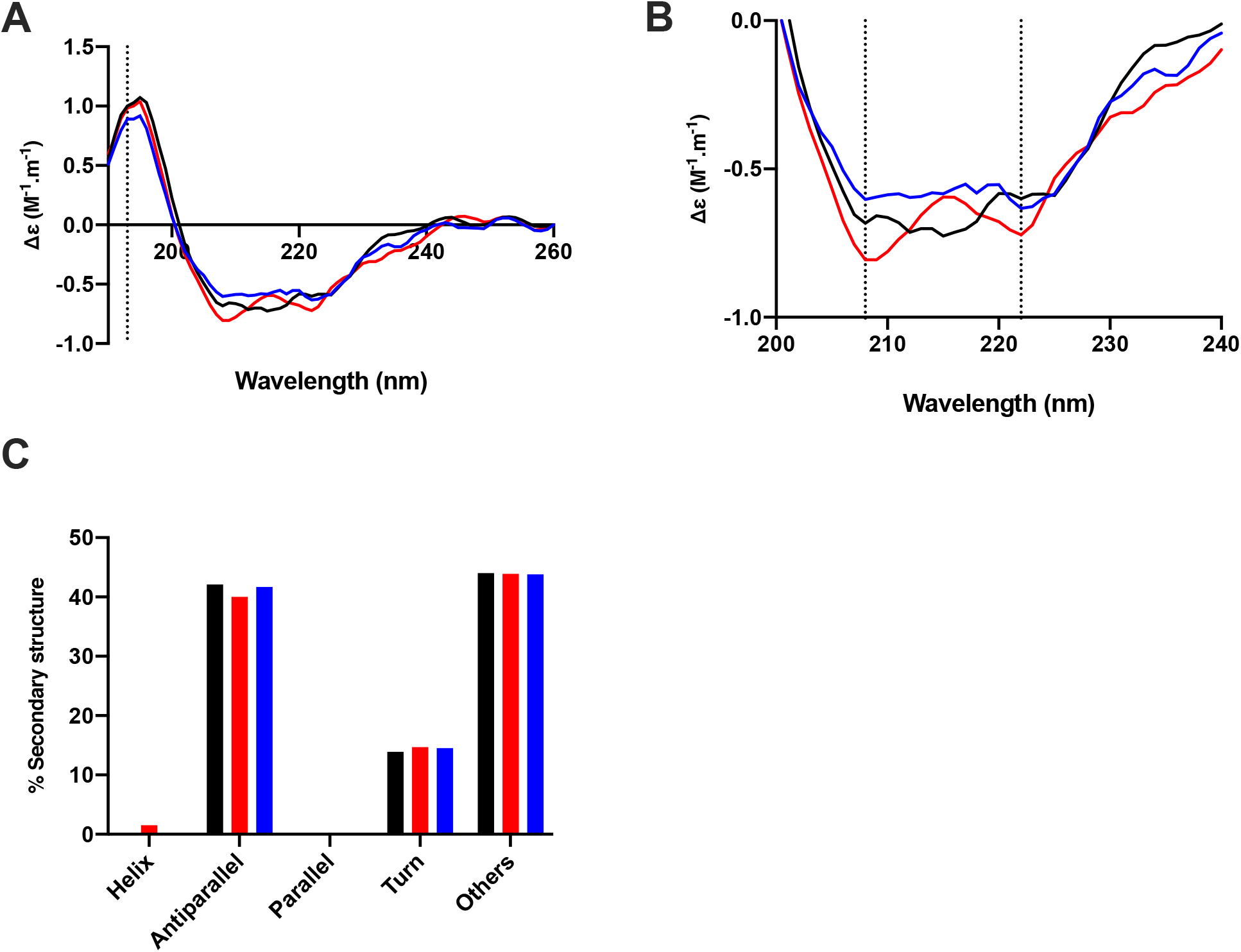
The conformational change of the SARS-CoV-2 S1 RBD observed in the presence of tinzaparin by circular dichroism (CD) spectroscopy. (**A**) Circular dichroism spectra (190 - 260 nm) of nCovS1RBD alone (black solid line), heparin (red solid line) and tinzaparin (blue) in PBS, pH 7.4. The dotted vertical line indicates 193 nm. (**B**) Details of the same spectra expanded between 200 and 240 nm. Vertical dotted lines indicate 222 nm and 208 nm. (**C**) Secondary structure content analysed using BeStSel for nCovS1RBD. α helical secondary structure is characterized by a positive band at ~193 nm and two negative bands at ~208 and ~222 nm (analysis using BeStSel was performed on smoothed data between 190 and 260 nm.3.2 Surface Plasmon Resonance binding studies.

**Figure 7:**
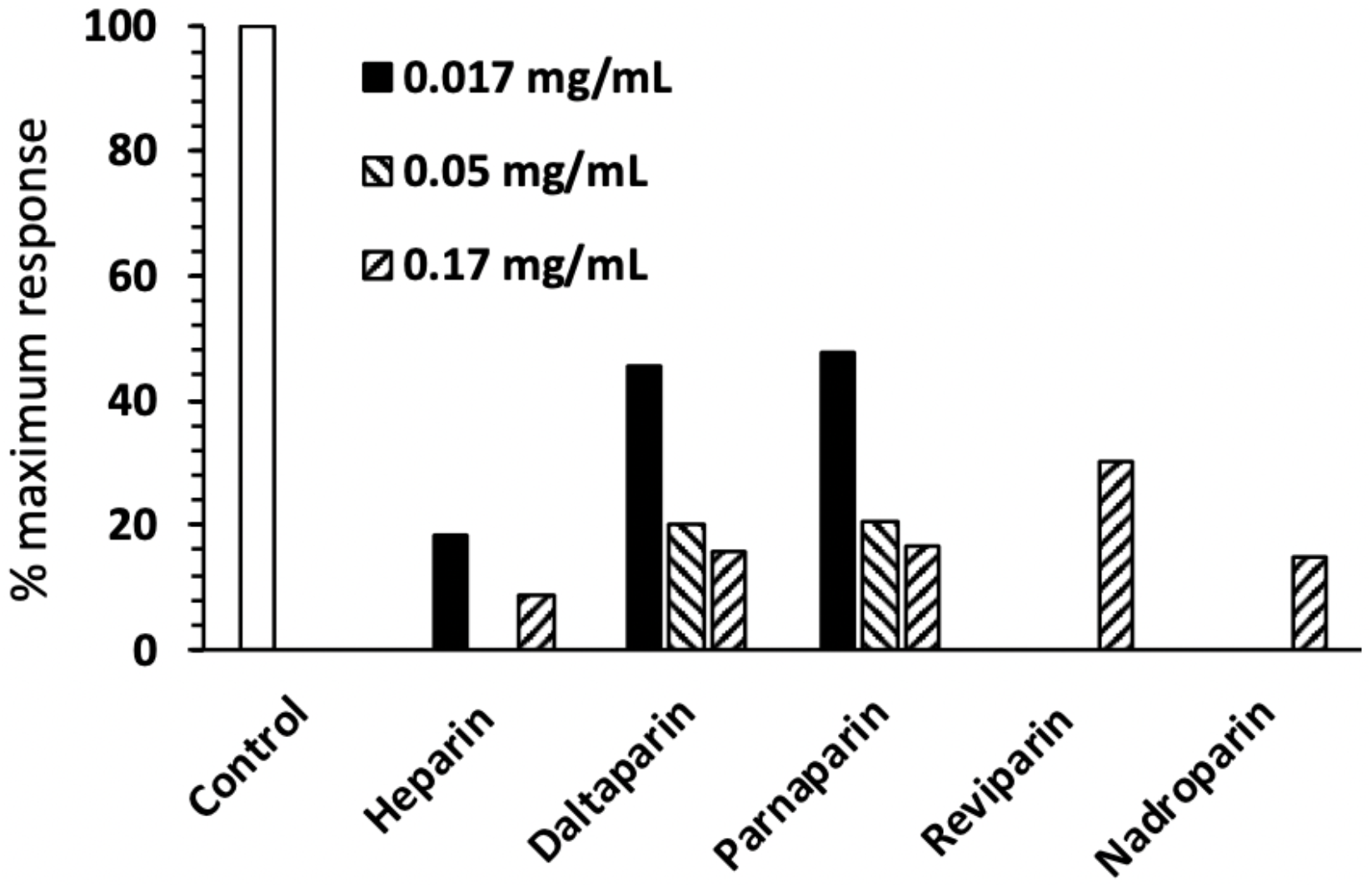
Competition of 800 nM SARS-cov-RBD binding to heparin by low molecular weight heparins. One mL SARS-CoV-2 S1 RBD (800 nM) was mixed with the appropriate low molecular weight heparin at the indicated concentration and injected over the biotinylated heparin sensing surface at 500 μl.min^−1^. Binding was measured after the surface had returned to PBST and expressed as a % of SARS-CoV-2 S1 RBD binding measured in the absence of competitor.

## Discussion and Conclusion

The rapid spread of SARS-CoV-2 represents a significant challenge to global health authorities and, as there are no currently approved drugs to treat, prevent and/or mitigate its effects, this makes repurposing existing drugs both a timely and appealing strategy. Unfractionated heparin and LMWHs are well-tolerated anticoagulant drugs, that have been used successfully for many years with limited and manageable side effects.

Studying SARS-CoV-2 Spike protein structure and behaviour in solution is a vital step for the development of effective therapeutics against SARS-CoV-2. Here, the ability of the SARS-CoV-2 S1 RBD to bind pharmaceutical LMWHs has been studied using spectroscopic techniques. The data show that SARS-CoV-2 S1 RBD binds to LMWHs and that upon binding, significant structural changes are induced.

The glycosaminoglycans class of carbohydrates are present on almost all mammalian cells and play a central role in the strategy employed by *coronaviridae* to attach to host cells. The GAG heparin has previously been shown to inhibit SARS-associated coronavirus cell invasion ^2,3,13^ and this, in concert with the LMWH data presented within this study, supports the use of GAG-derived pharmaceuticals as therapeutic agents against SARS-associated coronavirus. Furthermore, this study provides evidence for the repurposing of LMWHs as antiviral agents, providing a rapid countermeasure against the current SARS-CoV-2 outbreak.

It is noteworthy that even pharmaceutical-grade UFH and LMWH preparations remain a polydisperse mixture of natural products, containing both anticoagulant and nonanticoagulant saccharide structures. These may prove to be an invaluable resource for next-generation, biologically active, antiviral agents that display negligible anticoagulant potential, whilst the former remains tractable to facile, chemical (and enzymatic) engineering strategies to ablate their anticoagulation activities.

The further subfractionation of existing LMWH preparations against anticoagulant activities (with proven low-toxicity profiles, good bioavailability and industrial-scale manufacturing) for off-label pathologies, provides an attractive strategy for quickly and effectively responding to COVID-19 and for the development of next generation heparin-based therapeutics.

Such drugs will be amenable to routine parenteral administration through currently established routes and additionally, direct to the respiratory tract via nasal administration, using nebulised heparins, which would be unlikely to gain significant access to the circulation. Thus, the anticoagulant activity of UFH and LMWHs, which can in any event be engineered out, would not pose a problem. Importantly, such a route of administration would not only be suitable for prophylaxis, but also for patients under mechanical ventilation ^20^.

## Supplementary data

**Supplementary Figure 1:**
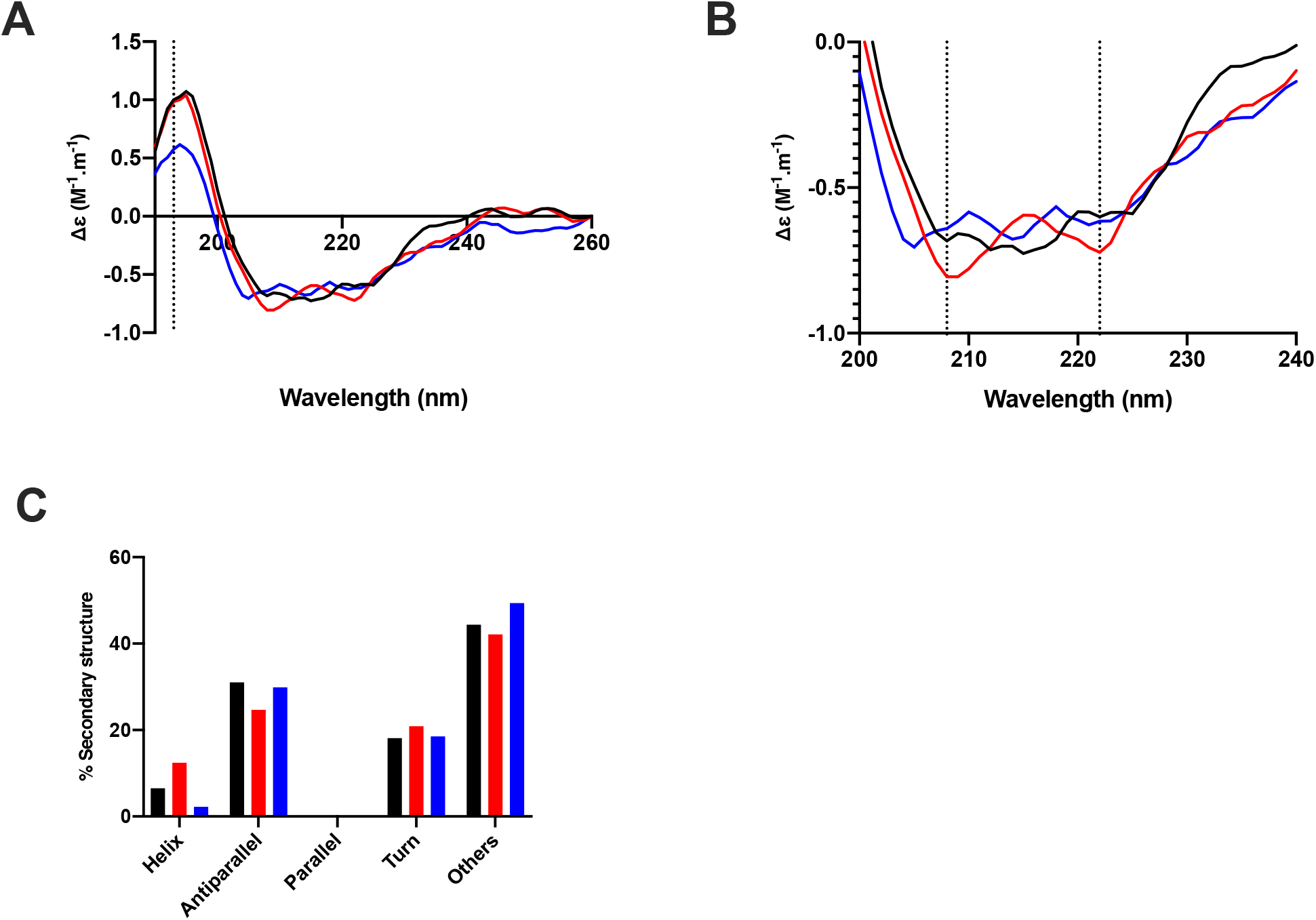
The conformational change of the SARS-CoV-2 S1 RBD observed in the presence of fondaparinux by circular dichroism (CD) spectroscopy. (**A**) Circular dichroism spectra (190 - 260 nm) of nCovS1RBD alone (black solid line), heparin (red solid line) and fondaparinux (blue) in PBS, pH 7.4. The dotted vertical line indicates 193 nm. (**B**) Details of the same spectra expanded between 200 and 240 nm. Vertical dotted lines indicate 222 nm and 208 nm. (**C**) Secondary structure content analysed using BeStSel for nCovS1RBD. α helical secondary structure is characterized by a positive band at ~193 nm and two negative bands at ~208 and ~222 nm (analysis using BeStSel was performed on smoothed data between 190 and 260 nm.

